# Glucagon-like peptide-1 receptor differentially controls mossy cell activity across the dentate gyrus longitudinal axis

**DOI:** 10.1101/2022.03.14.484169

**Authors:** Alex Steiner, Benjamin M. Owen, James P. Bauer, Leann Seanez, Sam Kwon, Jessica E. Biddinger, Ragan Huffman, Julio E. Ayala, William P. Nobis, Alan S. Lewis

## Abstract

Understanding the role of dentate gyrus (DG) mossy cells (MCs) in learning and memory has rapidly evolved due to increasingly precise methods for targeting MCs and for *in vivo* recording and activity manipulation in rodents. These studies have shown MCs are highly active *in vivo*, strongly remap to contextual manipulation, and that their inhibition or hyperactivation impairs pattern separation and location or context discrimination. What is not well understood is how MC activity is modulated by neurohormonal mechanisms, which might differentially control the participation of MCs in cognitive functions during discrete states, such as hunger or satiety. In this study, we demonstrate that glucagon-like peptide-1 (GLP-1), a neuropeptide produced in the gut and the brain that regulates food consumption and hippocampal-dependent mnemonic function, might regulate MC function through selective expression of its receptor, GLP-1R. RNA-seq demonstrated that most *Glp1r* in hippocampal principal neurons is expressed in MCs, and *in situ* hybridization revealed strong expression of *Glp1r* in hilar neurons. *Glp1r-ires-Cre* mice crossed with Ai14D reporter mice followed by co-labeling for the MC marker GluR2/3 revealed that almost all MCs in the ventral DG expressed *Glp1r* and that almost all *Glp1r*-expressing hilar neurons were MCs. However, only ~60% of dorsal DG MCs expressed *Glp1r*, and *Glp1r* was also expressed in small hilar neurons that were not MCs. Consistent with this expression pattern, peripheral administration of the GLP-1R agonist exendin-4 (5 μg/kg) increased cFos expression in ventral but not dorsal DG hilar neurons. Finally, whole-cell patch-clamp recordings from ventral MCs showed that bath application of exendin-4 (200 nM) depolarized MCs and increased action potential firing. Taken together, this study identifies a potential neurohormonal mechanism linking a critically important satiety signal with activity of MCs in the ventral DG that might have functional effects on learning and memory during distinct states.

## INTRODUCTION

The use of cutting-edge tools for recording and manipulating neurons has led to a rapid expansion in understanding the *in vivo* activity and functional roles of dentate gyrus (DG) mossy cells (MCs). Mossy cells are glutamatergic neurons with cell bodies in the DG hilus that provide both monosynaptic feedforward excitation and disynaptic feedforward inhibition of granule cells (Scharfman, 2016; Scharfman & Myers, 2012). Recording or imaging activity *in vivo* revealed that MCs are more active than granule cells, have multiple place fields, and strongly remap their spatial activity in response to contextual changes (Danielson et al., 2017; GoodSmith et al., 2017; GoodSmith et al., 2022; GoodSmith, Lee, Neunuebel, Song, & Knierim, 2019; Jung et al., 2019; Senzai & Buzsaki, 2017). Mossy cell ablation, inhibition, or excitation has shown their involvement in not only cognitive processes such as pattern separation and novelty detection that are critical for contextual and spatial memory (Bauer et al., 2021; Bui et al., 2018; Fredes et al., 2021; Jinde et al., 2012; X. Li et al., 2021), but also in a diverse repertoire of other behaviors and processes, including anxiety and avoidance (Botterill, Vinod, et al., 2021; Wang et al., 2021), neurogenesis (Oh et al., 2020; Yeh et al., 2018), and food intake (Azevedo et al., 2019).

Despite this strong foundation establishing that carefully tuned MC activity is necessary for aspects of episodic memory, a critical gap in understanding MC biology is how neuromodulatory systems regulate MC excitability. Elucidating networks such as neurohormonal signaling might clarify how MC circuits function during distinct motivational states. Aside from studies examining MC expression of monoamine (Etter & Krezel, 2014; Oh et al., 2020) and glucocorticoid (Patel & Bulloch, 2003) receptors, this endeavor has received limited attention.

Neurohormonal systems regulating feeding have consistently been shown to also regulate hippocampal function with consequences for learning and memory (reviewed in (Suarez, Noble, & Kanoski, 2019)). As such, we were intrigued that cell-specific RNA sequencing (RNA-seq) of hippocampal excitatory neurons revealed transcriptional enrichment of *Glp1r*, the gene encoding the glucagon-like peptide-1 receptor (GLP-1R) (Cembrowski, Wang, Sugino, Shields, & Spruston, 2016), in MCs. Other feeding-relevant endocrine receptors, including ghrelin, leptin, and insulin, were not reported as enriched in MCs. GLP-1 is produced in the distal gut and in the brainstem, where its role in the central regulation of feeding has been extensively described (Muller et al., 2019). Interestingly, GLP-1R signaling also promotes hippocampal-dependent spatial and associative learning (During et al., 2003; Isacson et al., 2011) as well as DG adult neurogenesis (Gault, Lennox, & Flatt, 2015; H. Li et al., 2010). However, the specific neuronal substrate within the hippocampus on which GLP-1 acts is not well defined. In this study, we characterize the expression and function of GLP-1R on MCs in the murine DG. We find that hippocampal GLP-1Rs are strongly and selectively expressed on ventral DG MCs and that ventral MC GLP-1Rs are functional both *ex vivo* and *in vivo*. Our results support future investigation of how ventral MC GLP-1Rs regulate cognitive and non-cognitive functions shown previously to be mediated by MCs.

## MATERIALS AND METHODS

### Animals

Male and female *Glp1r-ires-Cre* mice (RRID:IMSR_JAX:029283) (Williams et al., 2016) were crossed with Ai14D tdTomato reporter mice (RRID:IMSR_JAX:007914) (Madisen et al., 2010) to yield *Glp1r-ires-Cre* x Ai14D mice. All other mice used were male and female C57BL/6. Animals were group housed with a 12-hour light/dark cycle at 72 ± 2 °F with *ad libitum* access to food and water. All procedures were approved by the Vanderbilt Institute Animal Care and Use Committee.

### Drugs

Exendin-4 acetate was purchased from Cayman Chemical Company (Ann Arbor, MI).

### cFos expression following exendin-4 administration

Mice were habituated to the testing room for at least 1 hr, then administered exendin-4 (5 μg/kg in saline, s.c.) or saline, returned to their home cage, and transcardially perfused 90 mins later.

### Immunostaining

Terminal anesthesia, transcardial perfusion, and tissue sectioning were performed exactly as previously described (Bauer et al., 2021). For GluR2/3 immunostaining: two brain sections from each animal in each region of interest were selected. Dorsal DG slices were at approximately anterior/posterior (AP): −1.94 mm and ventral DG slices at approximately AP: −3.10 mm. Sections were permeabilized and blocked in 0.3% Tx-100 and 3% normal donkey serum (Jackson ImmunoResearch, West Grove, PA) in phosphate-buffered saline (PBS) for 2 hours. Sections were incubated overnight in rabbit anti-GluR2/3 (AB1506, Millipore Sigma, Burlington, MA, RRID:AB_90710, 1:200 dilution) at 4 °C. Sections were washed 3 x 10 mins in PBS, then incubated in donkey anti-rabbit Alexa 488 (Jackson ImmunoResearch, 1:1,000 dilution) at room temperature for 2 hrs. Sections were washed 3 x 10 mins in PBS, then incubated in 4′,6-diamidino-2-phenylindole (DAPI, Millipore, 1:5,000 dilution) in PBS at room temperature for 5 mins, washed in PBS, then mounted on slides using Fluoromount G (Electron Microscopy Sciences, Hatfield, PA). cFos immunostaining was performed identically, except permeabilization and blocking was done using 0.1% Tx-100, 1% normal donkey serum in PBS. Primary antibody was rabbit anti-cFos (226 003, Synaptic Systems, Goettingen, Germany, RRID:AB_2231974, 1:1,000 dilution), and secondary antibody was donkey anti-rabbit Alexa 488 (Jackson ImmunoResearch, 1:500 dilution).

### Microscopy and image quantification

Imaging was performed using an LSM 880 (Zeiss, White Plains, NY) equipped with a 20x Plan-Apochromat objective (NA = 0.8) and Zeiss Zen software for acquisition. All image analysis was performed blinded. For *Glp1r-ires-Cre* x Ai14D and GluR2/3 co-labeling, total numbers of tdTomato+ (i.e., Ai14D reporter), GluR2/3+, and tdTomato+GluR2/3+ neurons were counted within the hilus of each DG of each section and averaged across at least 2 sections per mouse. For cFos activation, the total number of cFos+ neurons within the DG hilus were counted and averaged across at least two sections per mouse. Images were processed using Fiji (Schindelin et al., 2012).

### Acute slice preparation

Acute brain slices were prepared from male and female juvenile (P19 to P35) C57BL/6 mice. Mice were decapitated under isoflurane, and their brains were removed quickly and placed in an ice-cold sucrose-rich slicing artificial cerebrospinal fluid (ACSF) containing (in mM): 85 NaCl, 2.5 KCl, 1.25 NaH_2_PO_4_, 25 NaHCO_3_, 75 sucrose, 25 glucose, 0.01 DL-APV, 100 kynurenate, 0.5 Na L-ascorbate, 0.5 CaCl_2_, and 4 MgCl_2_. Sucrose-ACSF was oxygenated and equilibrated with 95% O2/5% CO2. Horizontal slices (300-350 μm) were prepared using a vibratome (model VT1200S, Leica Biosystems). Slices were transferred to a holding chamber containing sucrose-ACSF warmed to 30°C and slowly returned to room temperature over the course of at least 30 min. Slices were then transferred to oxygenated ACSF at room temperature containing (in mM): 125 NaCl, 2.4 KCl, 1.2 NaH_2_PO_4_, 25 NaHCO_3_, 25 glucose, 2 CaCl_2_, and 1 MgCl_2_, and were maintained under these incubation conditions until recording.

### Electrophysiological recordings

Slices were transferred to a submerged recording chamber continuously perfused at 2.0 ml/min with oxygenated ACSF maintained at 30 °C. Putative hilar mossy cells were identified using infrared differential interference contrast on a microscope (Slicescope II, Scientifica) and recordings made as previously reported (Hedrick et al., 2017). Whole-cell patch-clamp recordings were performed using borosilicate glass micropipettes with tip resistance between 3 and 6 MΩ. Signals were acquired using an amplifier (Axon Multiclamp 700B, Molecular Devices). Data were sampled at 10 kHz and low-pass filtered at 10 kHz. Access resistances ranged between 16 and 22 MΩ and were continuously monitored before switching to current-clamp configuration. Changes greater than 20% from the initial value that were recorded at the end of an experiment were excluded from data analyses. Series resistance was uncompensated. Data were recorded and analyzed using pClamp 11 (Molecular Devices). Current clamp was performed using a potassium gluconate-based intracellular solution containing the following (in mM): 135 K-gluconate, 5 NaCl, 2 MgCl_2_, 10 HEPES (pH 7.0), 0.6 EGTA, 4 Na_2_ATP, and 0.4 Na_2_GTP, pH 7.3, at 281 mOsm). Input resistance was measured immediately after breaking into the cell and was determined from the peak voltage response to a 5 pA current injection. Following stabilization and measurement of the resting membrane potential, current was injected to hold all cells at a membrane potential between 60 – 65 mV, maintaining a common membrane potential that is within the reported resting membrane potential of hilar mossy cells to account for intercell variability. Hilar mossy cells were selected for recording based on the presence of a large, multipolar soma in the hilus. After achieving whole-cell configuration, hilar mossy cells were verified by a large whole-cell capacitance (>45 pF), a high frequency of sEPSCs (>5 Hz), and baseline action potential firing. Action potentials were characterized as having a mean amplitude of 83.988 ± 4.578 mV, a mean duration of 1.64 ± 0.09 ms, and a ratio of rising slope:decay slope greater than 2 (n = 7 cells from 6 animals), which is similar to that reported in the literature (Scharfman, 1992, 1995; Scharfman & Myers, 2012; Scharfman & Schwartzkroin, 1988).

### Statistical analysis

t tests (paired or unpaired, as appropriate) were used to compare two groups. To compare three or more groups, one- or two-way analysis of variance (ANOVA) as appropriate with Sidak’s post test was used. All tests were two-tailed. Analyses were performed using Prism 9 (GraphPad, San Diego, CA). Error bars depict standard error of the mean (SEM) unless otherwise noted.

## RESULTS

Using cell-specific RNA-seq, Cembrowski *et al*. previously reported that expression of *Glp1r* was enriched in MCs (Cembrowski et al., 2016). We further investigated this enrichment using HippoSeq, a publicly available tool to analyze the RNA-seq data generated by this study. *Glp1r* expression was only detectable in MCs and ventral CA3 pyramidal neurons (**Figure 1a**). Gene expression of the receptors for ghrelin, insulin, and leptin, which like GLP-1 are other feeding-relevant hormones previously shown to be important in hippocampal function (Suarez et al., 2019), was markedly lower than *Glp1r* expression and not enriched in MCs (**Figure 1b**). HippoSeq did not divide MCs into dorsal and ventral DG MCs, which is important because dorsal and ventral MCs differ not only molecularly (Blasco-Ibanez & Freund, 1997; Fujise, Liu, Hori, & Kosaka, 1998), physiologically (Bui et al., 2018; Fredes et al., 2021; Jinno, Ishizuka, & Kosaka, 2003), and anatomically (Botterill, Gerencer, Vinod, Alcantara-Gonzalez, & Scharfman, 2021; Houser, Peng, Wei, Huang, & Mody, 2020), but also in their role in cognitive function (Bauer et al., 2021; Botterill, Vinod, et al., 2021; Yassa & Stark, 2011). Therefore, to corroborate RNA-seq data and examine potential dorsal and ventral MC expression differences, we examined *in situ* hybridization (ISH) data for *Glp1r* from the Allen Mouse Brain Atlas (Lein et al., 2007) (**Figure 1c,d**). These data demonstrated *Glp1r* expression in neurons in both dorsal and ventral DG hilus, with markedly stronger staining in ventral DG, and limited expression elsewhere. Together, these data reveal that *Glp1r* in hippocampal excitatory neurons is enriched in MCs, and that *Glp1r* expression is strongest in hilar neurons of the ventral DG.

**Figure 1.**
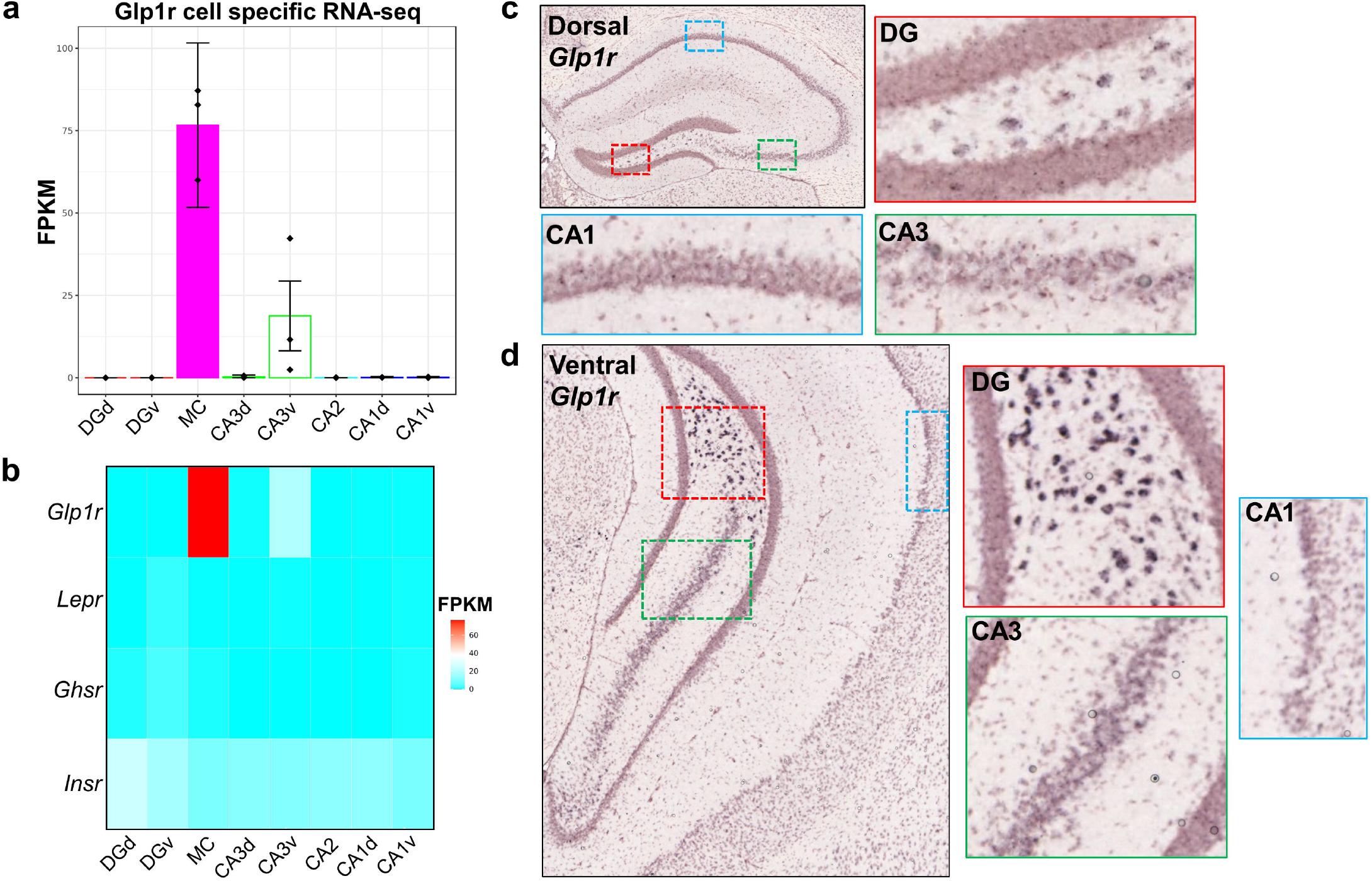
*Glp1r* gene expression in the hippocampus. **a,b)** HippoSeq, a publicly available mouse hippocampal principal neuron RNA-seq gene expression database, demonstrates that expression of *Glp1r*, the gene encoding GLP-1R, is highly enriched in MCs (a). Data in (a) are represented as mean ± 95% confidence intervals. Gene expression of other feeding-relevant hormone receptors in the hippocampus is much weaker than *Glp1r* expression and not enriched in MCs (b). *Lepr*, leptin receptor; *Ghsr*, growth hormone secretagogue receptor (ghrelin receptor); *Insr*, insulin receptor. FPKM, Fragments Per Kilobase of Exon Per Million Reads Mapped. **c,d)** *In situ* hybridization for *Glp1r* from Allen Mouse Brain Atlas corroborates RNA-seq data demonstrating most *Glp1r* expression is in hilar neurons consistent with MC expression, with strongest expression in the ventral DG hilus. There is limited expression in CA1 or CA3 pyramidal cell layers. From bregma, depicted dorsal DG is ~−1.9 mm and depicted ventral DG is ~−3.5 mm. Magnification factor is the same for all colored boxes.

Because HippoSeq only includes excitatory neurons and did not differentiate MCs between dorsal and ventral DG, the above RNA-seq and ISH data are unable to determine 1) whether all MCs express *Glp1r*, 2) whether all *Glp1r* is confined to MCs (since interneurons were not included in HippoSeq), and 3) whether these expression patterns differ between dorsal and ventral MCs. All of these might have important functional consequences. To approach these questions, we examined *Glp1r-ires-Cre* x Ai14D mice, which express the fluorescent reporter tdTomato in Cre-positive neurons (**Figure 2**). This genetic reporter approach differs from ISH in that it allows visualization of not only neuronal soma but also projections and permits colabeling with MC markers. Unlike ISH however, genetic reporter intensity cannot be used as a proxy for relative expression. In dorsal hippocampus, tdTomato was almost exclusively expressed in hilar cells, with almost no expression in the granule cell layer, area CA3, and area CA1. In ventral hippocampus, tdTomato was expressed in hilar cells, including at the extreme ventral pole. Expression was sparse in ventral CA3, though much more prevalent than in dorsal CA3. Ventral CA1, as in dorsal CA1, was largely devoid of tdTomato expression. In both dorsal and ventral sections, a strong band of tdTomato immunoreactivity was found in the inner molecular layer, the site of MC terminals. Finally, strong expression of tdTomato was found throughout the brain in blood vessels, consistent with known localization of GLP-1R in brain arteriolar smooth muscle and endothelial cells (Nizari et al., 2021).

**Figure 2.**
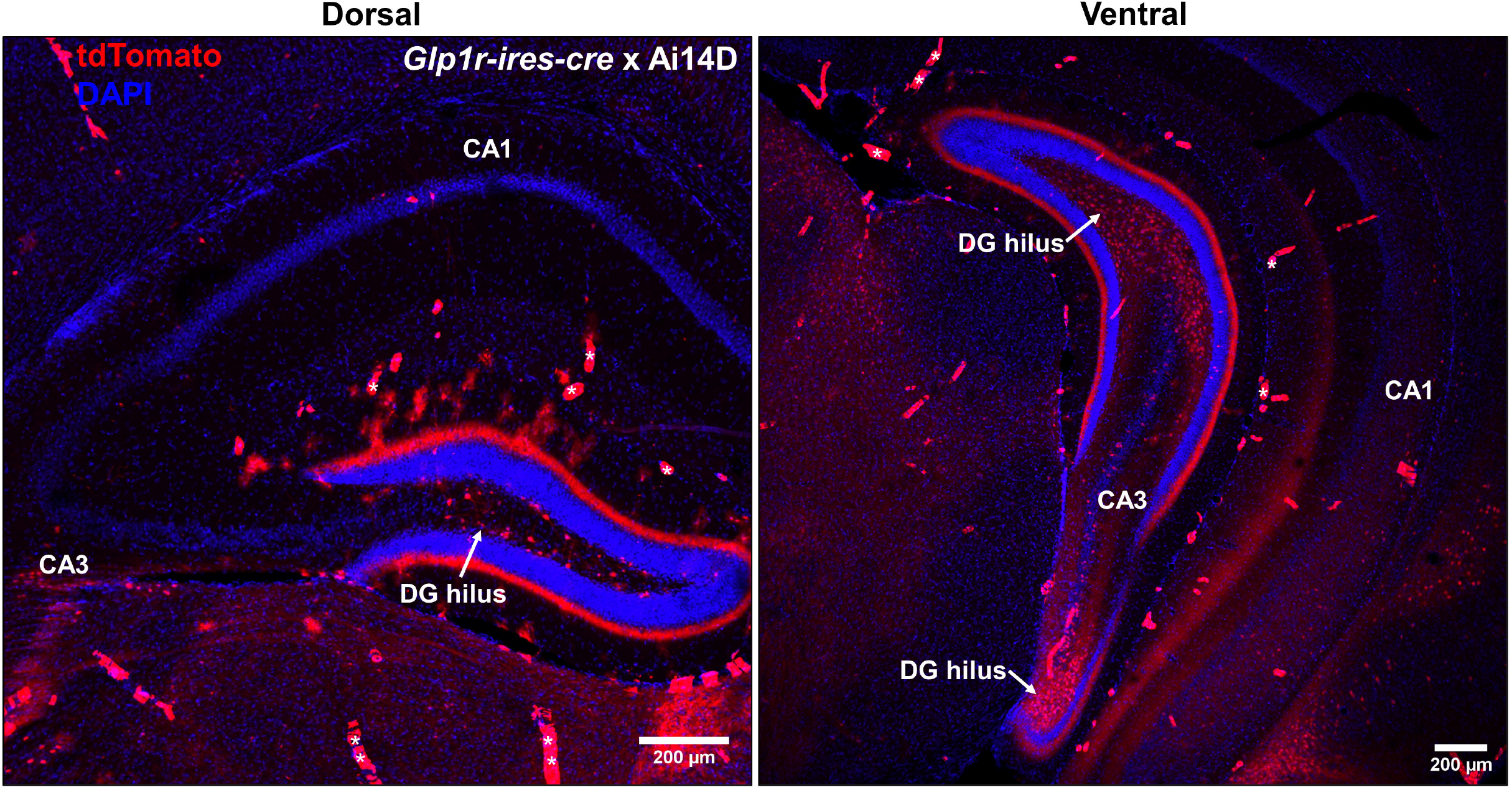
Genetic reporter for *Glp1r-ires-Cre* expression in dorsal and ventral hippocampal formation. Confocal microscopy of dorsal and ventral hippocampal sections from *Glp1r-ires-Cre* mice crossed with Ai14D genetic reporter line (*Glp1r-ires-Cre* x Ai14D), in which the red fluorescent protein tdTomato is expressed in Cre-positive neurons. Imaging revealed hilar expression in both dorsal and ventral hippocampus, along with sparse ventral CA3 expression. There was also strong tdTomato expression in the DG inner molecular layer, consistent with MC terminals. *Glp1r* is also strongly expressed in the brain vasculature, exemplified by structures marked by asterisks.

We next tested whether hilar neurons expressing the fluorescent genetic reporter were MCs by co-labeling with GluR2/3, a marker of hilar MCs in both dorsal and ventral DG (Jiao & Nadler, 2007; Leranth, Szeidemann, Hsu, & Buzsaki, 1996) (**Figure 3a**). These studies revealed that MC expression of *Glp1r* and *Glp1r* specificity for MCs differed between dorsal and ventral MCs (**Figure 3b,c**). In the ventral DG, 94.8% ± 2.9% of GluR2/3+ MCs expressed *Glp1r* and 93.8% ± 1.9% of *Glp1r*-expressing hilar neurons expressed GluR2/3. However, in the dorsal DG, only 61.3% ± 10.4% of dorsal DG GluR2/3+ MCs expressed *Glp1r*, and 66.0% ± 9.7% of *Glp1r*-expressing neurons also expressed GluR2/3+, suggesting about one-third of dorsal *Glp1r*-expressing neurons were not MCs. Altogether, RNA-seq, ISH, and genetic reporter strategies suggest that *Glp1r* expression overall is more prevalent in ventral hippocampus, where it is expressed in a sparse population of CA3 pyramidal neurons and expressed in essentially all ventral MCs, while *Glp1r* is not universally expressed in dorsal MCs and is also expressed in non-MC hilar neurons. Furthermore, dense tdTomato-positive terminals in the inner molecular layer throughout the DG dorsoventral axis suggests that *Glp1r*-positive MCs innervate granule cells across all hippocampal lamellae.

**Figure 3.**
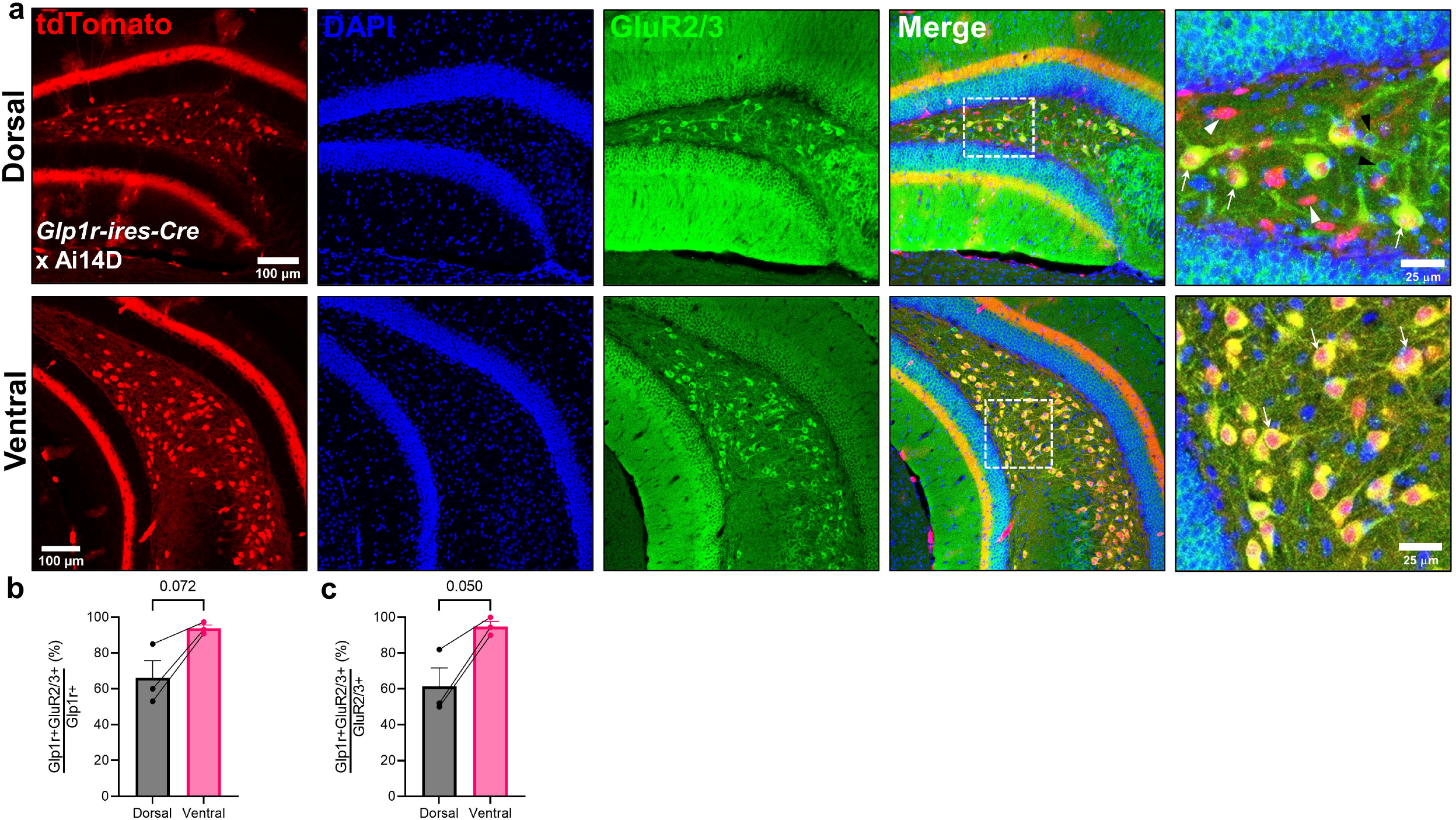
Hilar mossy cell *Glp1r* expression differs across the DG longitudinal axis. **a)** *Glp1r-ires-Cre* mice crossed with Ai14D reporter mice (*Glp1r-ires-Cre* x Ai14D) express tdTomato in Cre-positive neurons. Representative images from the dorsal and ventral DG are shown in the top and bottom rows of images, respectively, revealing tdTomato expression in hilar somata as well as a dense band of projections in the DG inner molecular layer. Sections were stained for GluR2/3, which in the hilus is a marker for MCs. Images on the far right are magnification of the boxed area. White arrowheads denote example *Glp1r*-positive/GluR2/3-negative neurons, black arrowheads denote example *Glp1r*-negative/GluR2/3-positive neurons, and white arrows denote example *Glp1r*-positive/GluR2/3-positive neurons. **b,c)** In the ventral DG, almost all MCs were *Glp1r*-positive, and almost all *Glp1r*-positive neurons were MCs. However, in the dorsal DG, only about 61% MCs were *Glp1r*-positive and 66% of *Glp1r*-expressing neurons were MCs. N = 3 mice. Paired t test, dorsal versus ventral: %*Glp1r*+GluR23+/*Glp1r*+: t(2) = 3.53, p = 0.072; % *Glp1r*+GluR23+/GluR23+: t(2) = 4.31, p = 0.050.

Glucagon-like peptide-1 and several GLP-1R agonists, including exendin-4, readily cross the blood-brain barrier (Kastin & Akerstrom, 2003; Kastin, Akerstrom, & Pan, 2002) where they have centrally mediated effects on cognition, feeding, and neurogenesis (During et al., 2003; Gault et al., 2015; Isacson et al., 2011; Kanoski, Fortin, Arnold, Grill, & Hayes, 2011). To investigate whether DG activity was changed by pharmacological activation of GLP-1Rs, we administered the GLP-1R agonist exendin-4 (5 μg/kg) peripherally, sacrificed the animal 90 mins later, and performed immunohistochemistry for the immediate early gene cFos. Consistent with stronger expression of *Glp1r* in ventral MCs, we found a significant increase in cFos following exendin-4 administration in the ventral but not dorsal DG (**Figure 4**).

**Figure 4.**
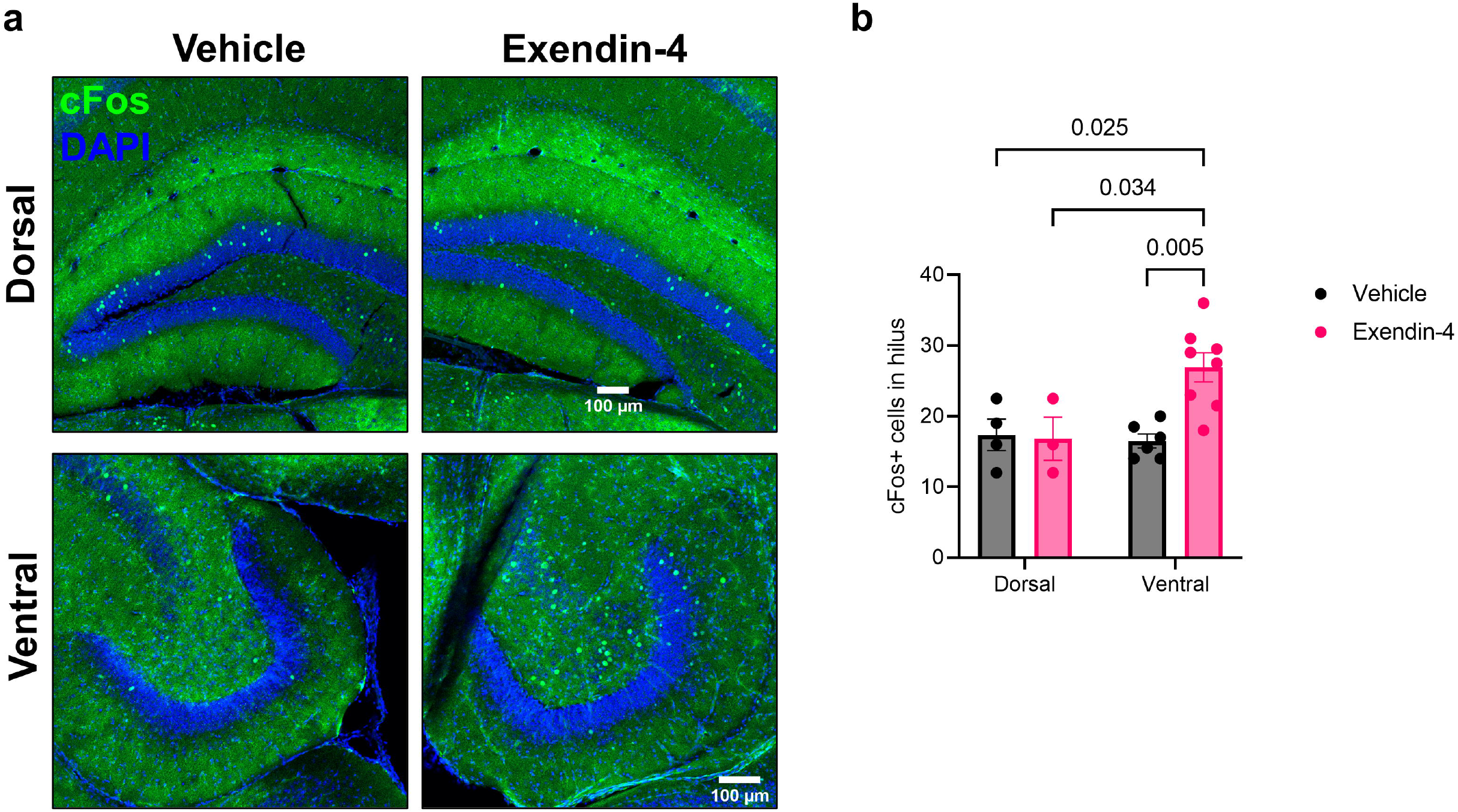
Peripheral administration of GLP-1R agonist increases ventral but not dorsal DG hilar cFos expression. Mice were administered the GLP-1R agonist exendin-4 (5 μg/kg) or vehicle and returned to their home cage. 90 mins later mice were sacrificed. Immunohistochemistry for the immediate early gene cFos revealed increased hilar cFos expression in ventral but not dorsal DG, quantified from horizontal and coronal sections, respectively. Two-way ANOVA: dorsoventral region x treatment interaction: F(1,17) = 6.154, p = 0.024; dorsoventral region: F(1,17) = 4.348, p = 0.052, treatment: F(1,17) = 4.999, p = 0.039. Pairwise comparison p values from Sidak’s multiple comparison tests are shown in the figure.

Along with *Glp1r* expression data, these findings suggest that GLP-1R activation might act directly on MCs to increase their firing. However, cFos cannot differentiate this possibility from network effects, such as activation of MCs and hilar GABAergic interneurons via GLP-1R-mediated activation of CA3c neurons and back projections to these DG hilar cell types, a process known to be particularly strong in the ventral hippocampus (Scharfman, 2007). To quantify the effects of GLP-1R activation on MC physiology, we performed whole-cell current-clamp recordings from ventral DG MCs to examine the effect of bath application of exendin-4 (200 nM) (**Figure 5a**), a dose used recently to characterize neuronal responses to GLP-1R activation *ex vivo* (Povysheva, Zheng, & Rinaman, 2021). We specifically examined ventral MCs because our expression data demonstrated that almost all ventral hilar neurons expressing *Glp1r* are MCs (**Figure 3**), which, in addition to our electrophysiological criteria (see methods), contributes to the likelihood that we were indeed recording from MCs. Bath application of 200 nM exendin-4 significantly increased the rate of spontaneous action potential firing (**Figure 5b-d**) compared to baseline in all cells. This was associated with a small but significant membrane depolarization compared to baseline (**Figure 5e,f**) and is consistent with actions of GLP-1 agonists in other neuronal populations (Cork et al., 2015; Povysheva et al., 2021).

**Figure 5.**
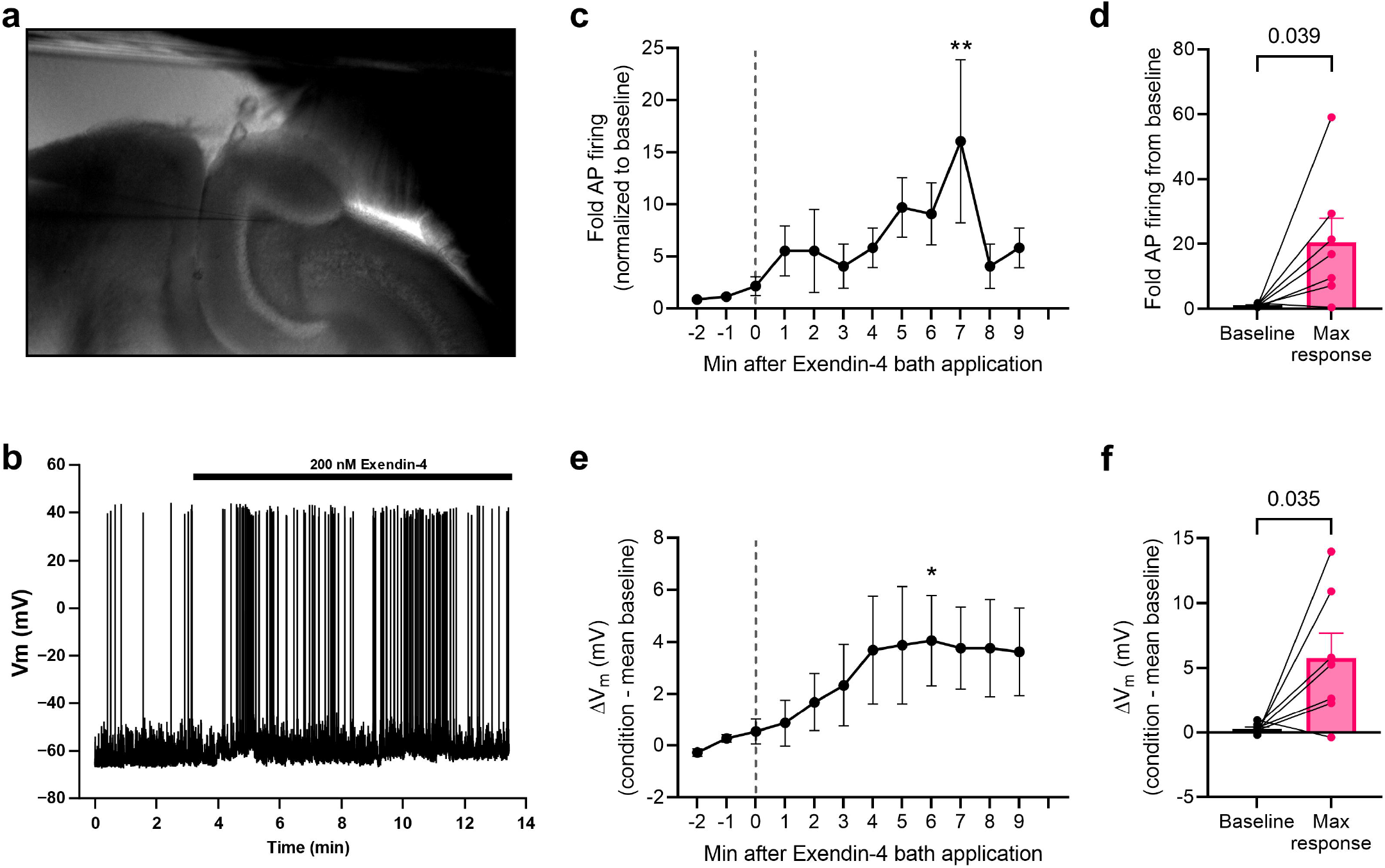
GLP-1R agonist depolarizes hilar MCs and increases action potential firing. **a)** Image of recording electrode in the hilus from a horizontal ventral DG slice. **b)** Representative current clamp recording of hilar MC following bath application of GLP-1R agonist exendin-4 (200 nM) showing increase in action potential firing and membrane depolarization. **c)** Mean action potential (AP) firing following exendin-4 (200 nM) bath application reveals overall increase in firing. Data are normalized to mean firing during the 2 min period prior to onset of bath application. One-way repeated measures ANOVA: F(11,66) = 2.194, p = 0.025. **p < 0.01 versus −1 min baseline (Sidak’s multiple comparison test). N = 7 cells from 6 mice. **d)** Maximal firing rate in the 9 min period following exendin-4 bath application is significantly increased from baseline. Paired t test versus −1 min baseline: t(6) = 2.62, p = 0.039. **e)**Change in membrane potential (V_m_) also increased significantly following exendin-4 bath application. One-way repeated measures ANOVA: F(11,66) = 3.112, p = 0.0020. *p < 0.05 versus −1 min baseline (Sidak’s multiple comparison test). N = 7 cells from 6 mice. **f)**Maximal V_m_ change in the 9 min period following Exendin-4 bath application is significantly increased from baseline. Paired t test versus - 1 min baseline: t(6) =2.71, p = 0.035.

## DISCUSSION

Identifying non-synaptic mechanisms that regulate MC activity is critically important to bridge our current understanding of MC function, primarily gleaned from activity manipulation studies, with MC function during specific physiological and behavioral states. In this study, we demonstrate that GLP-1 signaling via GLP-1Rs on MCs may be an important neurohormonal mechanism regulating MC activity, as well as a cellular target for GLP-1 analogues already in clinical use. Using RNA-seq, ISH, and genetic reporter lines, we found that 1) ventral MCs strongly express *Glp1r*, 2) that GLP-1R agonist depolarizes MC membrane potential and increases action potential firing *ex vivo*, and 3) that peripheral administration of GLP-1R agonist increases ventral DG hilar neuron activity *in vivo*. Interestingly, dorsal MCs differed markedly from ventral MCs in their expression of *Glp1r*, where it is expressed in only about two-thirds of MCs, and only about 60% of *Glp1r*-positive hilar neurons were MCs. *Glp1r* expression as revealed by ISH in dorsal hilar neurons was substantially weaker than in ventral hilus, which may contribute to our inability to detect cFos activation in the dorsal hilus by peripheral GLP-1R agonist. Alternatively, dorsal hilar *Glp1r*-positive neurons that were not MCs may play an active role in inhibiting local hilar neurons following GLP-1R agonist administration, also accounting for an absence of detectable cFos increase.

As is the case for the hippocampal formation as a whole (Fanselow & Dong, 2010; Strange, Witter, Lein, & Moser, 2014), an appreciation of meaningful differences between dorsal and ventral DG MCs continues to evolve. These include marked differences in protein expression (Blasco-Ibanez & Freund, 1997; Cembrowski et al., 2016; Fujise et al., 1998), activity (Bui et al., 2018; Fredes et al., 2021; Jinno et al., 2003), connectivity (Botterill, Gerencer, et al., 2021; Houser et al., 2020), and effects on cognitive and behavioral function (Bauer et al., 2021; Botterill, Vinod, et al., 2021; Yassa & Stark, 2011). For instance, relevant to our *Glp1r* expression findings, in mice the calcium binding protein calretinin is strongly and selectively expressed in MCs in the ventral and intermediate DG whereas most dorsal MCs are calretinin-negative (Blasco-Ibanez & Freund, 1997; Fujise et al., 1998). Ventral MCs show markedly greater intrinsic bursting than dorsal MCs due to differential expression of persistent sodium currents (Jinno et al., 2003). Finally, ventral MCs are significantly more active in novel contexts and this novelty detection can gate contextual fear conditioning whereas dorsal MC activity plays a less specific role in this form of learning (Fredes & Shigemoto, 2021; Fredes et al., 2021). Thus, our findings that ventral MCs and the ventral DG are activated by GLP-1R agonist may have several behavioral consequences. Whether GLP-1R signaling improves or degrades performance in DG-dependent cognition is likely task-dependent and difficult to predict. For example, either optogenetic inhibition (Bui et al., 2018) or chemogenetic excitation (Bauer et al., 2021) of ventral MCs impairs spatial encoding during an object location memory task, suggesting that disruption of distinct activity bidirectionally has a degradative effect on encoding. However, GLP-1R signaling may enhance contextual fear conditioning in familiar environments (Fredes et al., 2021) and perhaps restore MC activity that is necessary for mnemonic function in pathological conditions in which MCs are lost, such as epilepsy (Blümcke et al., 2000; Bui et al., 2018).

It is important to note that MC protein expression may be highly species dependent, which has previously been appreciated in calretinin staining between mice, rats, and monkeys (Blasco-Ibanez & Freund, 1997; Gulyás, Miettinen, Jacobowitz, & Freund, 1992; Miettinen, Gulyás, Baimbridge, Jacobowitz, & Freund, 1992; Seress, Nitsch, & Leranth, 1993). Along these lines, recent work in rats identified *Glp1r* expression in ventral CA1 neurons and showed that GLP-1R signaling through these neurons had functional effects on food intake and operant responding (Hsu, Hahn, Konanur, Lam, & Kanoski, 2015; Hsu et al., 2018). In contrast, murine RNA-seq, ISH, and a *Glp1r-ires-Cre* x Ai14D reporter line cross did not demonstrate appreciable ventral CA1 *Glp1r* expression (**Figures 1 and 2**). Even within species, different transgenic approaches to visualize neuronal *Glp1r* show notable differences. For example, the *Glp1r-ires-Cre* knockin line crossed with Ai14D mice shown in the present study exhibits similar dorsal and ventral DG *hilar* neuronal expression to a *Glp1r-Cre* BAC transgenic line (Cork et al., 2015; Richards et al., 2014) and a GLP-1R-mApple BAC transgenic line (Graham et al., 2020). However, these three lines differ in their reporter expression in other hippocampal cell types, including DG granule cells and pyramidal neurons in CA1 and CA3, which contrast with RNA-seq and ISH data. That three different transgenic lines demonstrate hilar neuronal expression adds confidence to the expression of *Glp1r* in MCs. However, variable expression elsewhere suggests a need for caution when making claims as to the degree of *Glp1r* expression in these other hippocampal fields.

Our findings set the stage for several avenues of future study for understanding healthy brain function and for the development of therapeutic interventions. Basic motivational states, such as hunger or thirst, interact with cognitive processes to guide adaptive behavior. For instance, in a state of hunger, an adaptive response is to seek and consume food. This process is facilitated by pairing the hungry state with prioritized recall of food-paired contexts. Indeed, such fundamental motivational states are coded in hippocampal firing (P. J. Kennedy & Shapiro, 2004; Pamela J. Kennedy & Shapiro, 2009; Wood, Dudchenko, Robitsek, & Eichenbaum, 2000), but how state is communicated to the hippocampus is not well understood. As a satiety signal (Hsu et al., 2018; Muller et al., 2019; Trapp & Richards, 2013), GLP-1 signaling is one potential mechanism that might couple satiety or absence of satiety with distinct MC function. Indeed, because GLP-1Rs are G-protein-coupled receptors, even relatively short-lived prandial increases in central GLP-1 levels might be well situated to have long-lasting effects on MCs that far outlast the presence of the hormone itself. Supporting this notion, MCs were recently shown to be more active in the fed than fasted state (Azevedo et al., 2019). Finally, the unique vulnerability of MCs in disorders such as epilepsy and the cognitive consequences of their loss coupled with the current widespread clinical use of GLP-1R pharmacotherapies encourages translational investigation into whether targeting MC GLP-1Rs can preserve or enhance pattern separation function to facilitate episodic memory in neurological or psychiatric disease.

## Acknowledgements

This work was supported by National Institutes of Health (NIH) Grant MH116339 (ASL), the Nicholas Hobbs Discovery Grant (ASL), and the Vanderbilt Faculty Research Scholars award (WPN). Experiments and data analysis were performed in part using the Vanderbilt University Medical Center Cell Imaging Shared Resource (supported by NIH grants CA68485, DK20593, DK58404, DK59637, EY08126, and S10 OD021630). The authors have no conflicts of interest to declare.

## Data availability statement

The data that support the findings of this study are available from the corresponding author upon reasonable request.

